# SPART, a versatile and standardized data exchange format for species partition information

**DOI:** 10.1101/2021.03.22.435428

**Authors:** Aurélien Miralles, Jacques Ducasse, Sophie Brouillet, Tomas Flouri, Tomochika Fujisawa, Paschalia Kapli, L. Lacey Knowles, Sangeeta Kumari, Alexandros Stamatakis, Jeet Sukumaran, Sarah Lutteropp, Miguel Vences, Nicolas Puillandre

## Abstract

A wide range of data types can be used to delimit species and various computer-based tools dedicated to this task are now available. Although these formalized approaches have significantly contributed to increase the objectivity of SD under different assumptions, they are not routinely used by alpha-taxonomists. One obvious shortcoming is the lack of interoperability among the various independently developed SD programs. Given the frequent incongruences between species partitions inferred by different SD approaches, researchers applying these methods often seek to compare these alternative species partitions to evaluate the robustness of the species boundaries. This procedure is excessively time consuming at present, and the lack of a standard format for species partitions is a major obstacle. Here we propose a standardized format, SPART, to enable compatibility between different SD tools exporting or importing partitions. This format reports the partitions and describes, for each of them, the assignment of individuals to the “inferred species”. The syntax also allows to optionally report support values, as well as original trees and the full command lines used in the respective SD analyses. Two variants of this format are proposed, overall using the same terminology but presenting the data either optimized for human readability (matricial SPART) or in a format in which each partition forms a separate block (SPART.XML). ABGD, DELINEATE, GMYC, PTP and TR2 have already been adapted to output SPART files and a new version of LIMES has been developed to import, export, merge and split them.

## Introduction

Species delimitation (SD) is a burgeoning, fully fledged research field in systematic biology (Sites & Marshall 2003; Camargo & Sites 2013; Flot 2015, Ducasse et al. 2020). SD benefits from the interpretation of species as independent evolutionary lineages (De Queiroz 1998, 2007) that can be distinguished from each other using a variety of operational SD criteria (Samadi & Barberousse 2006). In integrative taxonomy (Dayrat 2005; Padial et al. 2010), various lines of evidence and a wide range of data types can be used in formalised analytical workflows to propose species hypotheses, from DNA barcodes to phylogenomic data, discrete morphological characters, morphometric measurements, ecological traits, geographic occurrence, bioacoustic signals, metabolomic profiles, and others (Miralles et al. 2020).

If many, and among them the earliest, formalised SD procedures are mostly carried out manually, e.g. by comparing trees with the geographic occurrence of individuals, calculating correlations between geographic and genetic distances, assessing steepness of hybrid zones, or seeking for correlation between genetic distance and morphological characters (Good & Wake 1992, Wiens & Penkrot 2002, Vieites et al. 2009, Flot et al. 2010, Weisrock et al. 2010, Puillandre et al. 2012a, Miralles & Vences 2013, Derkarabetian & Hedin 2014, Dufresnes et al. 2015), a substantial number of computer-based tools has been developed to delimit species, often based on statistical criteria. These programs can analyse large datasets, with a strong focus on the use of sequence data (Table 1). These methods have significantly contributed to increase the objectivity, repeatability, and speed of species delimitation inferences under different mathematical models and assumptions (e.g. Multispecies coalescent model, DNA barcode gap, haplotype fields of recombination, cf. de Queiroz 1998, 2007, Knowles & Carstens 2007, Yang & Rannala 2010, Carstens et al. 2013, Leavitt et al. 2015, Rannala 2015).

**Table 1.**
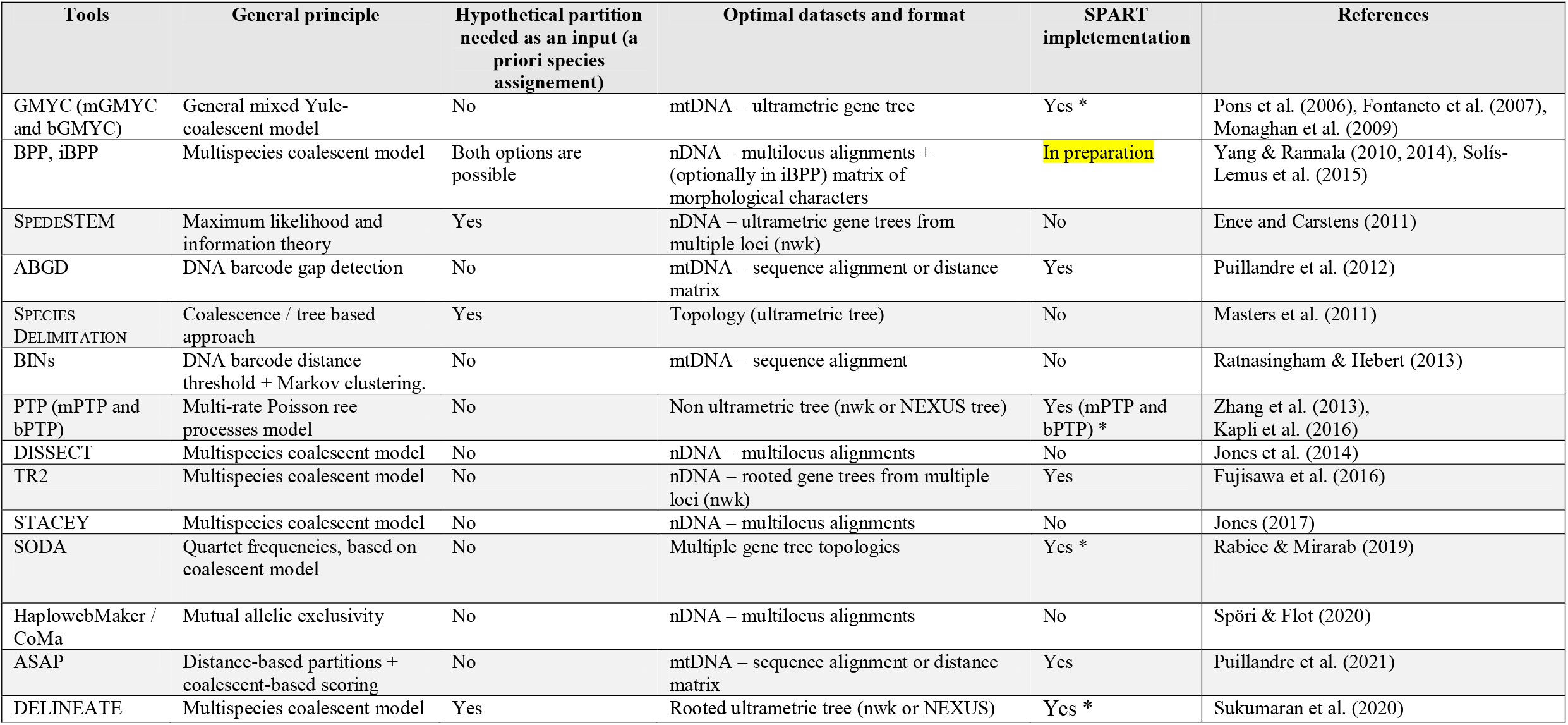
Automatedtools dedicated to species delimitation. Abbreviations used: mtDNA, mitochondrial DNA; nDNA, nuclear DNA. Note that for programs marked with an asterisk (GMYC, PTP, SODA, DELINEATE) GUI-driven versions with SPART implementation have been prepared in the context of the iTaxoTools project but SPART output is not yet provided by all available versions. Other programs (ABGD, ASAP, TR2) already include native SPART output.

Although the number and importance of SD tools is likely to sharply increase in the immediate future, they are not yet routinely used in the majority of alpha-taxonomic studies that result in the naming of over 15,000 new species of organisms every year (Miralles et al. 2020). One obvious shortcoming is the lack of interoperability among the various independently developed SD programs, and the lack of comprehensive software suites that offer various user-friendly features, such as those for data visualization and comparison of results across methods. For instance, incongruent species partitions resulting from different SD approaches applied to a given dataset are common. They can even be significant, if not striking in some cases (such as excessive splitting or lumping leading to highly different number of species delimited; Carstens et al. 2013, Miralles & Vences 2013, Dellicour & Flot 2015, Kapli et al. 2016, Postaire et al. 2016, Renner et al. 2017 for empirical cases; and Sukumaran & Knowles 2017, Chan et al 2020, Luo et al. 2018, Mason et al. 2020 and Zhang et al. 2011 for more methodological studies on SD limitations). Integrative taxonomists will seek to compare these alternative species partitions across SD approaches (but see Rannala 2015), and eventually estimate their robustness by integrating other data sources (morphological variation, geographic distribution, etc), in order to make an informed choice–a procedure that is excessively time consuming at present, given the lack of a standard format for species partitions.

The main output of species delimitation, and therefore of any SD program, is a species partition. The term “partition” here follows the set theory concept: the organization of a set of *elements* into mutually-exclusive and jointly-comprehensive *subsets*, not including the empty subset (Hrbacek & Jech 1999). In an SD application, the *elements are individuals* (i.e. samples or specimens), and a specific species delimitation hypothesis is a particular assignment (i.e. a *partition*) of these individuals to subsets, where *each subset corresponds to a distinct inferred species*. Categories resulting from an SD analysis have been referred to by various terms, such as primary species hypothesis, operational taxonomic unit (OTUs), barcode index number (BINs; Ratnasingham & Hebert 2013), or even cluster (without any particular status (Fig. 1)), but all of them match the aforementioned definition of a subset. Furthermore, while some tools produce *de novo* species partitions (i.e. directly aggregating individuals into species hypotheses; exploratory methods), others statistically compare competing species hypotheses that have been defined *a priori* (hypothesis-testing methods), and these programs require a species partition as input. SD methods may also assign scores, either to the entire inferred partition (e.g., ASAP-score in the program ASAP; Puillandre et al. 2021), to the distinctiveness of each subset from the others (e.g., posterior probabilities in the programs BPP and bPTP; Yang & Rannala 2010; Zhang et al. 2013), or to the presence of each individual in a given subset (e.g., probability of placement in calculation of BINs, Ratnasingham & Hebert 2013).

**Figure 1.**
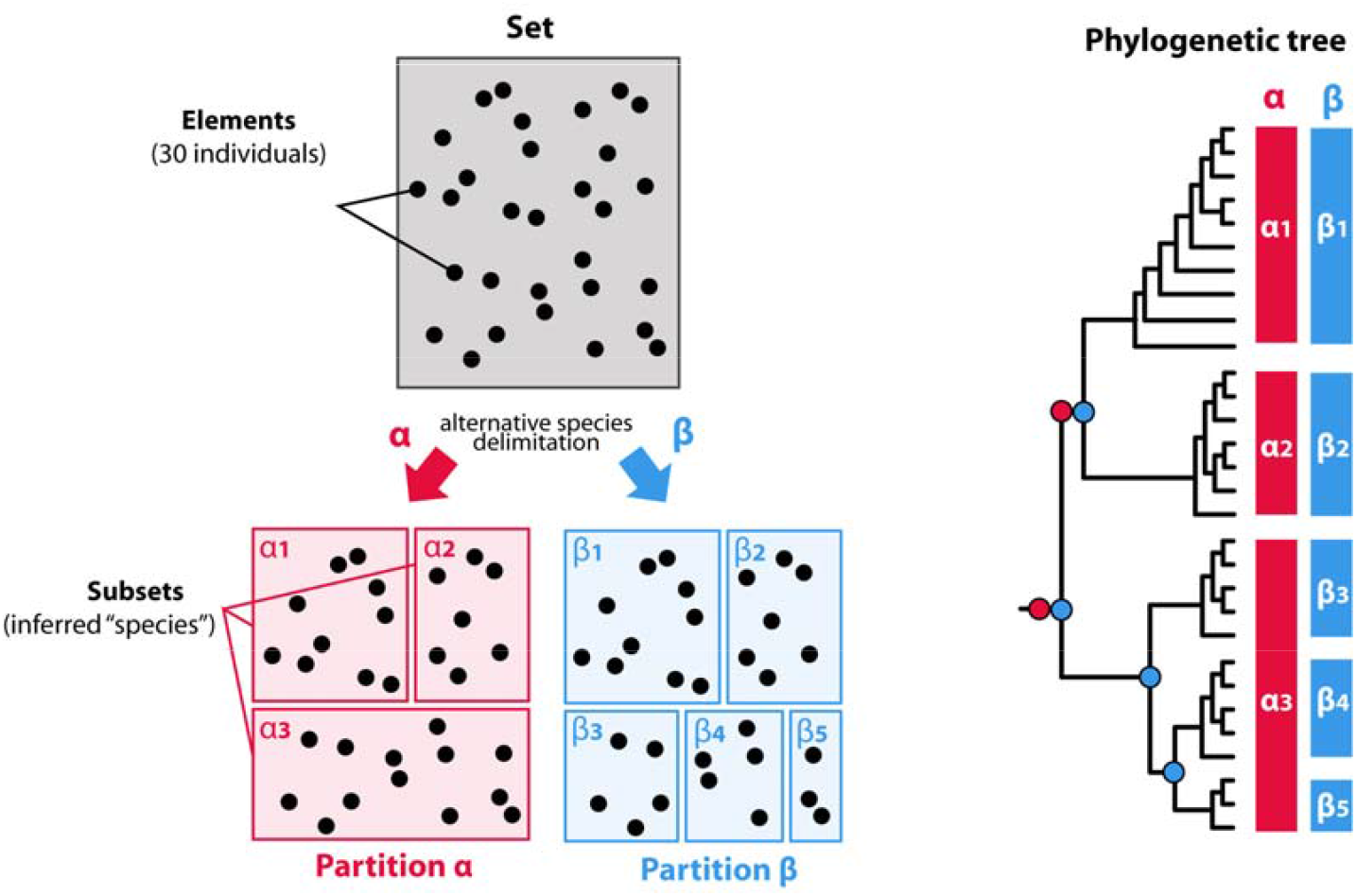
In mathematics, a partition of a set is a grouping of its elements into non-empty subsets, in such a way that every element is included in exactly one such subset. The main output of a species delimitation inference therefore corresponds to a partition, independently of the theoretical context, the biological input data, or the algorithms/models used. In our example, a set of 30 specimens is split by two different methods into two alternative partitions α and β, corresponding to 3 and 5 putative species (subsets), respectively. For the sake of clarity, these two alternative species partitions are represented as boxes reported next to each “species clade” in a phylogenetic tree, with hypothetical speciation events highlighted by circles via a corresponding color. Note that not all SD methods rely on a tree topology, and may therefore delimit non-monophyletic units (e.g., methods based on morphological or molecular divergence).

### A standardized Species PARTition format (SPART)

Typically, each SD program exports the resulting species partitions in its own idiosyncratic format. Some, for instance, provide a table of assignments of individual specimens to the subsets (e.g. GMYC) while others, conversely, list the different subsets with the included individuals (e.g. ABGD, PTP), whereas again others graphically report subsets on a tree topology (e.g. GMYC). These different formats may or may not include complementary data (e.g., scores, topologies, metadata, number of species delimited, etc.), and are not designed to be parsed by other tools for downstream analyses. Their manual conversion into a versatile and easily reusable plain text species partition (e.g., CSV) is not always straightforward. It can be particularly error prone and time consuming with large datasets, as species delimitations on several hundreds, or even thousands, of specimens are becoming common practice in molecular taxonomy (e.g., Ahrens et al. 2016, Renner et al 2017, Garcià-Melo et al. 2019, Hoffmann et al. 2019, Solihah et al. 2020, Christodoulou et al. 2020).

We here propose a standardized species partition format, SPART, to enable compatibility between different tools producing (export) or using (import) species partitions. Our format facilitates:

1. statistical comparison of different alternative species partitions such as their overall congruence, similarity or resolving power, identification of the subsets that are congruently delimited (currently implemented in the program LIMES v2.0; Ducasse et al. 2020);
2. assessment of multiple competing SD hypotheses, including those used as input in e.g. DELINEATE and BPP to evaluate them (Sukumaran et al. 2020, Yang & Rannala 2010);
3. visualization and comparison of species partitions (e.g., DNA-based species partitions compared with manually-edited species partitions obtained from alternative methods and datasuch as Principal Component Analysis of morphometry, haplotype networks, geographic distribution, habitat type, external phenetic similarity, or simply, current taxonomy);
4. extraction, from original data files, of specific data for each subset under different species partition assumptions (e.g. lists of molecular and morphological diagnostic character states, descriptive statistics characterizing each of the inferred species, or ecological or distributional traits); and
5. potential taxonomic reassignment of specimens in databases.

More generally, the SPART format is designed to be versatile and fully integrative in the sense that it can include any species partition descriptors, independently of the method or data-type used to generate the species partition (Fig. 2). SPART does not convey any interpretation on the quality of the species partition, nor on the pros and cons of the methods used to define them, but is simply a common format that seeks at homogenising the way species partitions are recorded. It can therefore be implemented in any method used to generate one or several species partitions as output. Likewise, any method using (analysing, comparing, automatically reassigning or graphically representing) multiple subsets of specimens might benefit from being able to import SPART files as input data.

**Figure 2.**
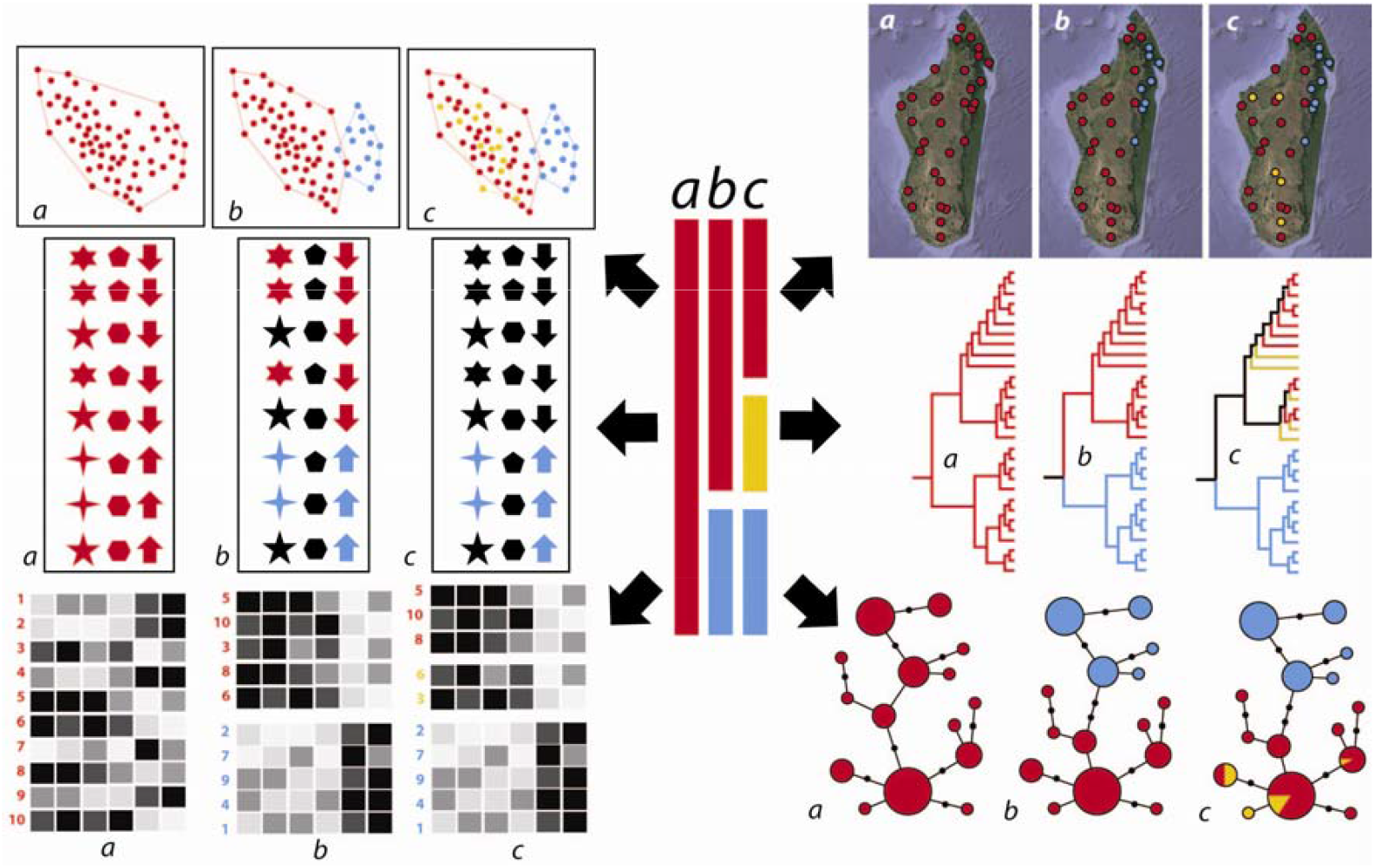
Illustration of exemplary potential applications of a species partition (SPART) file. If it can be parsed by other programs, SPART might facilitate the exploration of taxonomic datasets under various delimitation assumptions (such as morphometric Principal Component Analysis, automated extraction of diagnostic traits, heatmap of meristic morphological traits, distribution map, mitochondrial DNA-based phylogenetic tree, or haplotype network from nuclear DNA). In the present example, the partition b represents the optimal delimitation from a taxonomic perspective.

**Figure 3.**
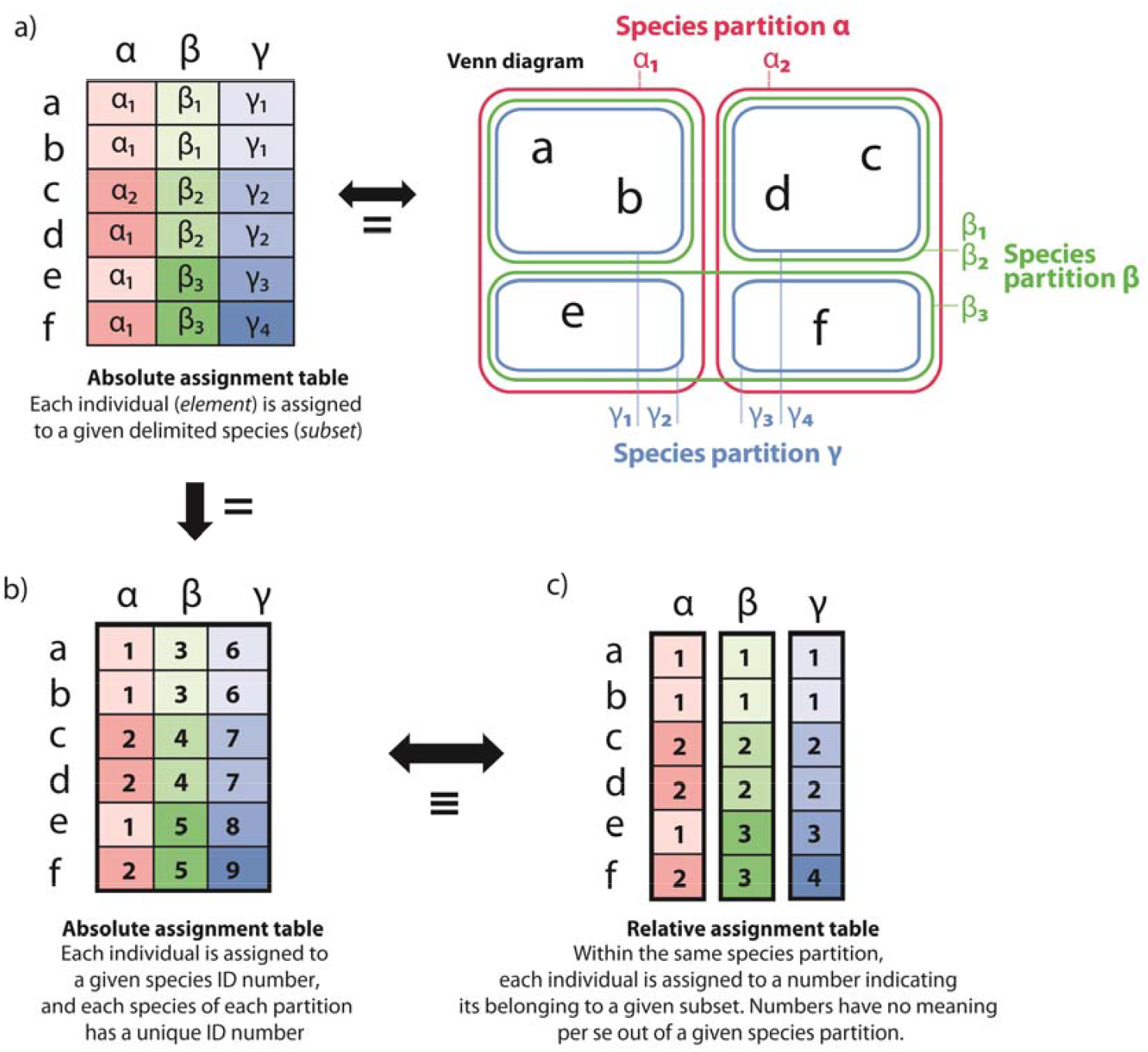
The SPART format can combine alternative species partitions of a same set of individuals (elements) into a unique multiple species partition file. (a) Example of set comprising six individuals split by three distinct SD analyses, resulting in three distinct species partitions (α, β and γ). All these species partitions are hierarchically compatible (i.e. they conform to the mathematical definition of nested sets), with the exception of the pair α - β (Venn diagram representing the alternative species partitions on the right, and corresponding assignment table on the left). These alternative species partitions can be coded in SPART either (b) by using a unique numbering for all the three species partitions (so that each species partition has its own set of species (subset) numbers) or (c) by using one numbering system per species partition. The latter representation allows combining different species partitions into a multiple species partition file without having to adjust each species or cluster number (subset). Both (b) and (c) are fully equivalent in SPART format, because the coding of each partition is independent from the others (subset assignment numbers have no meaning *per se*, they only indicate, within each partition, the common assignment to a specific subset).

### Matricial and serial implementation of the SPART format

SPART files include information on one or multiple species partitions for a given set of elements (i.e. individuals) and use standardized terminology to denote the number of species partitions included in the file (“N_spartitions”) and for each partition, the number of individuals (“N_individuals”), number of subsets (“N_subsets”), and the assignment of individuals to subsets (“Assignment”) (Supporting information 1). The syntax also allows to optionally include support values for species partitions, subsets, and the assignment of individuals to subsets, as well as original trees and the full command line used in the respective SD analyses, the program version number as well as comments and species partition comparison indices as calculated with LIMES 2.0, a new version of LIMES (Ducasse et al. 2020) recently published.

To account for the diversity of possible future applications, we propose two variants of the SPART format (for details see Supporting information 1). Both of these use largely the same terminology but represent the data differently:

The first SPART variant is optimized for human readability and its syntax has been designed to be compatible with Nexus (a widely used data format in phylogenetic inference software: Maddison et al. 1997). This allows to include SPART specifications as blocks in Nexus files if required by future applications. If information from multiple partitions is included, then it is combined into a single block, presenting the respective assignments and assignment scores per individual from different species partitions concatenated on a single line, separated by separator symbols. This enables easy manual transformation into a spreadsheet format if required. Due to the presentation of information from multiple partitions in one block as a concatenated matrix, we denote this variant as *matricial SPART* format, or simply SPART.

The second SPART variant is optimized for machine readability, and relies on XML (eXtensible Markup Language), a lightweight data-interchange format that can be easily parsed and written by software tools, while it can still be read and written by humans as well. When information from multiple partitions is included, each partition forms a separate block containing information on the number of subsets, individual assignments and assignment scores. We therefore denote this variant as *SPART*.*XML* format.

### Tools already implementing SPART and future perspectives

The proposed format is already implemented in several widely-used SD programs. Both the matricial SPART and SPART.XML output files are already generated by GUI-driven standalone versions (https://github.com/iTaxoTools) of ABGD, ASAP, GMYC, PTP, mPTP, TR2 and DELINEATE (Vences et al. submitted), by the native Python version of TR2, and in the web versions of ABGD and ASAP; and in progress for the Python versions of GMYC and PTP. Furthermore, the species partition comparison tool LIMES v2.0 has been expanded to import, export and convert SPART files, in particular to (1) compare, by calculating indices (e.g., *Ctax, Ratx, Match Ratio*, cf. Ducasse et al. 2020) for species partitions from SPART files (including each one or several species partitions); (2) merge species partitions included in different SPART files into one SPART file, (3) import species partition(s) table(s) from spreadsheet editors such as Microsoft EXCEL and save it (them) into a single SPART file. A new software tool named SPARTMAPPER has also been developed; it takes SPART files as input along with a tab-delimited series of geographical coordinates linked to specimen names, plots the distribution of alternative delimited species on a map, and exports a .kml file to visualize this information in Google Earth.

<u>[note to reviewers: all of these new software versions will be released before or upon publication of this manuscript, and the web based ABGD and ASAP are already freely available since February 2021</u> (https://bioinfo.mnhn.fr/abi/public/abgd/abgdweb.html, https://bioinfo.mnhn.fr/abi/public/asap/); <u>for review, various preliminary Windows executables are available under this link:</u> https://hidrive.ionos.com/share/ohaymwcgjd#$/Win%20executables]

In the context of future work, we envisage the development of visualization tools to automatically illustrate information from species partitions along with support values and phylogenetic hypotheses (Fig. 1). There is still a long way to go before programs will be able to infer species based on combining evidence using different data sources such as genetics, morphology, ecology, behaviour, geographic distribution, etc. However, eventually, reliable computer-based, species delimitation procedures that mirror the procedures of integrative taxonomy will be at the core of next generation taxonomy (Vences 2020). Our SPART data exchange format would thus contribute to this next generation taxonomy, by simplifying computational approaches to completing the inventory of life on Earth.

## Acknowledgments

We are grateful to Susanne Renner who stimulated this work by leading the priority program SPP 1991 “Taxon-Omics” of the Deutsche Forschungsgemeinschaft (DFG), specifically in the context of a grant on taxonomic data integration and management (RE 603/29-1), and to many members of the Taxon-Omics consortium and Guillaume Achaz (MNHN) for fruitful discussion. MV and SK were supported by DFG grant VE247/20-1, NP by the European Research Council (ERC) under the European Union’s Horizon 2020 research and innovation programme (grant agreement No. 865101), and AS and SL by the Klaus-Tschira foundation.

