## Appendix for "SPART, a versatile and standardized data exchange format for species partition information"

**Appendix 1: Technical description of the Spart format (species partition)**

(version 12/03/2021)

1. Background, environment and software implementation	page 2

2. Terminology	page 4

3. Technical details	page 5

4. Format description: matricial SPART (Nexus-like)	page 5

5. Format description: SPART.XML	page 12

**Participants in the initial development of the spart format:**

Brouillet S. (programming for ABGD, ASAP and other tools of the future MNHN platform),

Ducasse J. (programming for LIMES),

Kumari S. (programming for iTaxoTool which will implement various SD methods)

Miralles, A (design, MNHN),

Puillandre N. (design, MNHN),

Vences, M. (design, TU Braunschweig)

**1. Background, e****nvironment and software implementation**

The spart formats

Two implementations of the spart format are proposed:

► a matricial spart format (SPART) in which for each individual (sample), multiple species partition assignment is included in a concatenated, table-like format. This format has been designed to be intuitively understandable by humans, facilitating manual editing and import into table editors, and has a syntax largely compatible with the nexus format, commonly used in phylogenetics, thus facilitating its inclusion as a separate block into nexus files if required by future analysis software.

► a SPART.XML format in which the information for each species partition is provided in a separate block, and in which blocks are serially appended one after each other. This format is optimized for being machine-readable and its syntax follows the XML language.

We strongly recommend using the extensions *".spart*" and "*spart.xml*" for the matricial and XML implementation, respectively.

Current implementation (march 2021):

Programs currently implementing SPART : ABGD, ASAP, GMYC, PTP, SODA, TR2, DELINEATE, LIMES.

LIMES v2.0 (link to be added) will act as a central platform for converting and modifying spart files.

LIMES v2.0 is compatible with matricial spart (.spart) files; spart.xml compatibility will be implemented in the next version, together with the possibility to convert between matricial SPART and SPART.XML files. In addition, standalone and web-based tools will be implemented in the future to easily convert spart from and to spart.xml files.

LIMES v2.0 can read one or several species partitions from a CSV formatted document (thus including manually created spartitions), merge species partitions (= spartitions) from several single and/or multiple spartitions files, extract them, and export them into a single multi-spartitions spart file.

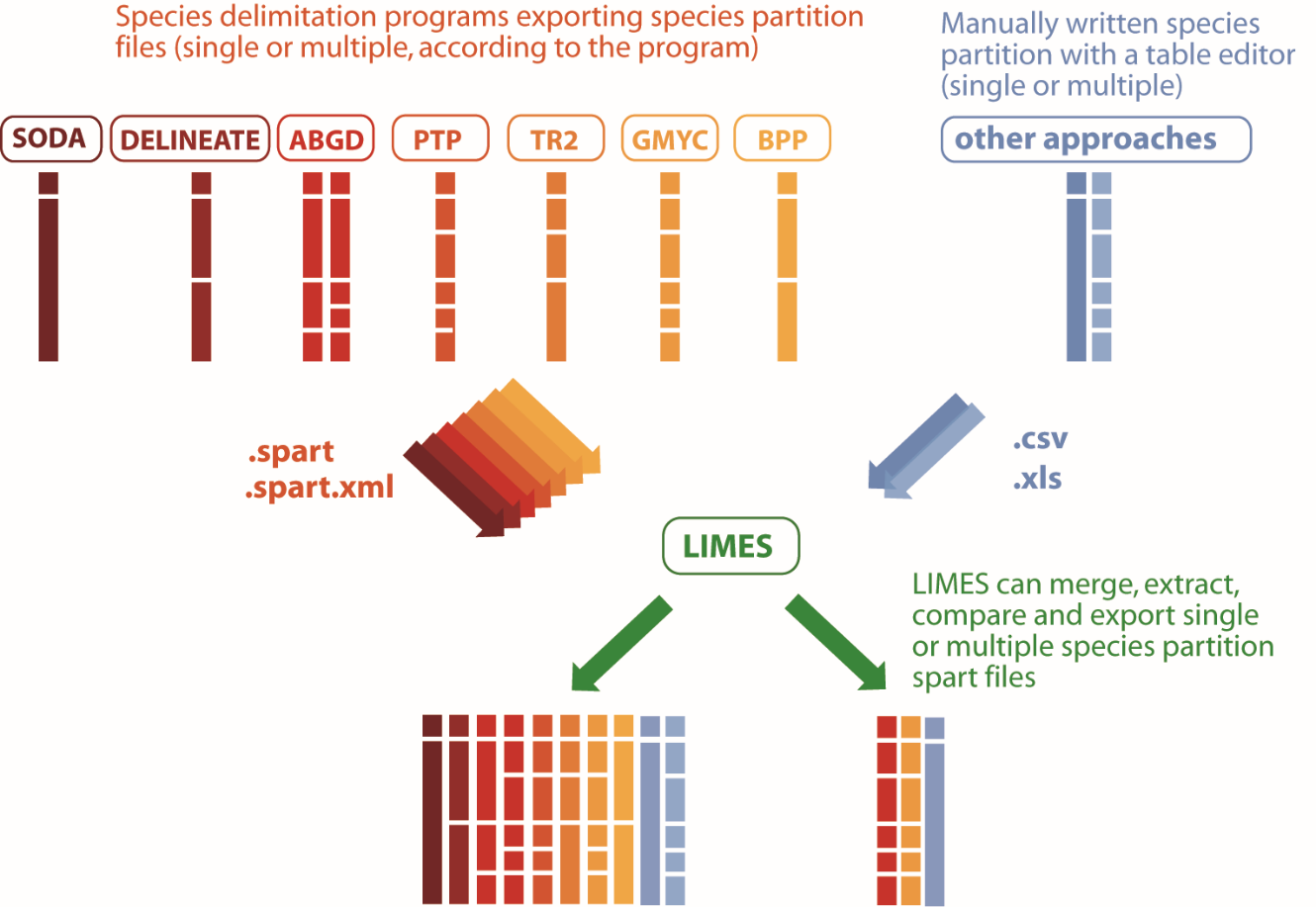

**2. Terminology**

**Species partition (Spartition):** Distribution (classification or assignment) of all individuals into multiple subsets, according to a given method.

**Subset:** elementary unit of the results (of the partition); usually a “species”, but can also be defined as a cluster or an operational taxonomic unit (OTU) or a molecular operational taxonomic unit (MOTU) or a barcode index number (BIN), a population (e.g. STRUCTURE) or any other kind of unit, depending on the computational analysis performed.

**Individual:** elementary unit of the dataset; usually equals a sample or a specimen in the SD analysis which in most cases will represent an individual organism (but can also be for instance an isolate/culture in microbiology).

**Spartition score:** any score attributed to the species partition as a whole (one score per species partition, usually corresponding to one score per SD analysis).

**Subset score:** any score attributed to each subset (to quantify its distinctiveness relative to the others).

**Individual score:** any score attributed to the assignment to a subset proposed for each individual.

Note: inclusion of these three scores in a spart file is optional, but if any such scores are calculated by an SD software, we recommend that the output spart files should include this information in the proposed format.

**Single species partition file:** Usually, the result of a species delimitation analysis (SD). Most often, each SD program is expected to export a file with a single species partition, but exceptions exists (e.g., ABGD typically provides several partitions and thus may either export multiple single species partition files, or one multiple species partition file).

**Multiple species partition file:** the information of several single species partitions merged into a single file, either as direct output from some SD programs, or by merging single species partition files using LIMES (or other tools with this functionality). The respective developers of most SD programs will typically implement the export of a single species partition file per analysis.

Note: ABGD and ASAP are already producing multiple species partitions as a result of a single analysis. So ideally, the user should be allowed to select which species partitions to export at the end of the analysis, and whether this should be done as multiple single species partition files, or as a single multiple species partition file. Typically, both ABGD and ASAP will provide a list of partitions, but some of them are often unrealistic (especially for ABGD, when they are far from the barcode gap), and the user may not desire to include them in the spart file exported by the program and used for further analysis.

**Block**: To enable compatibility with programs using Nexus as input file, the matricial spart file is conceived as a single block. Its start is indicated by an initial line specifying "begin spart;" and its end is indicated by a line specifying "end;".

In the spart.XML format, all information of one species partition is provided as one block separate from other blocks (species partitions).

**Command (matricial SPART format):** A command corresponds to a section/field intended to provide a specific type of information or instruction (e. g. “N_spartitions”, “N_subsets”, “N_individuals”, “Individual_assignment”, including the subsequently given details and values, each correspond to a different command).

**Command title:** Specific title given to a given command (e. g. “N_spartitions”, “N_subsets”, “N_individuals”, “Individual_assignment”).

**4. Format description: matricial spart**

**1) Character set :**

**By default:** All the 95 ASCII printable characters are allowed in the entire format (incl. space)

!"#$%&'()*+,-./0123456789:;<=>?@ABCDEFGHIJKLMN

OPQRSTUVWXYZ[\]^_`abcdefghijklmnopqrstuvwxyz{|}

**Exception for individual (sample) names (assignment list):** only numbers, capital and lower case letter and underscore (no spaces nor any other diacritic signs) are allowed: 0123456789ABCDEFGHIJKLMNOPQRSTUVWXYZ_abcdefghijklmnopqrstuvwxyz

We recommend to use underscore to replace any forbidden character

Any individual name including another character should lead to a specific error message

**Exception for partition names (ex. in N**_**spartitions):**

These six characters are strictly prohibited :

~~comma~~ ( , )

~~colon~~ ( : )

~~slash~~ ( / )

~~semi colon~~ ( ; )

~~opening and closing brackets~~ ( [ )	( ] )

Any partition name including one of these six characters should lead to a specific error message.

More generally, we recommend to only use numbers, capital and lower case letters (i.e. to use underscores to replace any other symbols), as above (individual samples).

**Exception for species assignment (Individual_assignment) :** only positive integers are allowed (0,1, 2, 55, 101, 102, etc.)

Any assignment using another character should lead to a specific error message

**2) Scores**

Scores will often represent a proportion (between 0 and 1) such as posterior probabilities or bootstrap proportions, but can also take very small values (e.g., likelihood scores).

Spartition, species and individual **scores** are either in the form of fixed-point notation (e.g. 0.093) or in the form of floating point numbers (scientific (exponential) notation (e.g., 9.30E-02)).

Negative values are permitted (e.g. log likelihood values)

**Question marks** (?) represent missing data.

A score category can be removed if (and only if) totally empty: in particular (but not limited to) cases where there is no spartition score at all, or no subset score at all, or no individual score at all.

**3) Separators**

Spart should be robust against whitespaces (we prefer to not take any risk, although Nexus separates e.g. taxon names and characters by whitespaces). The spart format avoids using tabs or spaces as separators. **The only separators allowed are the following:**

**Colons (:)** first order separator (although the term separator is not fully appropriate here). Colons are used to separate an item followed by a list of attributes sharing the same order.

**Slashes (/)** second order separator, used to separate values corresponding to different species partitions.

**Comma (,)** third order separator, used only for partition scores and subset scores.

**Equality signs (=),** separating a Command title from the values presented after the sign.

Additional “pseudo-separators”:

**End of line:** to end every line (within a given command) referring to an individual sample. Each new command (section) starts by a new line. End of line are NOT allowed within an individual sample line (ex. with a line of “Individual_assignement).

It is important that programs reading matricial spart files accept all possible end of lines, i.e., Windows (CR-LF), Unix (LF) and old Mac (CR)

**Semi-colon:** to indicate the end of a command. Only the semi-colon indicates “end of the command”. That means that it can appear after the last line, or in the line afterwards. The two following examples are equivalent and correct:

| **N**_**spartitions =** 3:      CO1_ABGD, 0.98 **/**              test_BPP, 0.95 **/**              PCA_phenotype,?  ; | **N**_**spartitions =** 3:      CO1_ABGD, 0.98 **/**              test_BPP, 0.95 **/**              PCA_phenotype,?; |
| --- | --- |

**Brackets** [in order to isolate a comment like in this sentence].

A comment can appear embedded within a command (between a command title (beginning)

and the end (;)). As brackets are used to frame a comment, a comment cannot contain a bracket embedded within it. *Exemple: [this comment is ~~[not]~~ correct]*

**4) Syntax**

**Spaces**: should have no influence (Spart should be robust against whitespaces).

For example : N_individuals = 5 / 5, N_individuals =5/5 , and N_individuals =5/ 5 should all be correct and equivalent (i.e software should be able to read a file written by hand and containing minor errors like these above).

Nevertheless, we recommend to implement (to automatically generate

e.g. as output of SD programs) only the following format: **N_individuals = 5 / 5**

**Commands** (N_spartitions, N_subsets, N_individuals, Individual_assignment…): are not case sensitive and the following examples all are equivalent and should be readable by programs: N_SUBSETS, n_subsets or N_SUBsets

We recommend to implement (to automatically

generate) only the following format: **N_subsets**

**Command order:**

The order of the compulsory commands must be respected:

1: Project_name, 2: Date, 3: N_spartitions, 4: N_individuals, 5: N_subsets, 6: Individual_assignment

The optional commands must appear after the compulsory

commands, but their respective order is free.

**Begin and end of spart file (spart block):**

Because in the matricial spart format, all information resides in one block, the beginning of the block is specified at the very beginning of the file (i.e., before the Project_name command):

begin spart;

and the end is specified either at the very end of the file (either after the last compulsory command if only compulsory commands are included in the file; or after the last optional command):

end;

Commands in the matricial spart format (SPART), exemplified by a multiple partition file

(but note that most SD programs will usually export a single species partition file)

| **Compulsory commands** | ***These commands need to be present in any spart file.***  *If any of them is missing, some programs using the spart file will possibly not work or output error messages.-> ERROR MESSAGE* |
| --- | --- |
| **begin spart;** | *Starting line of the spart file. Indicates the begin of a block (the entire spart file is conceived as a single block).* |
| **Project**_**name =** my_three_delimitations**;** | *Name given for this new project* |
| **Date =** 2020-09-21T07:26:10+00:00**;** | *Date and time (standard ISO 8601 recommended) in which* ***this*** *specific spart file was generated. The date is mandatory but the format is flexible.*  *These three examples are correct :*  Date=2021-03-04T16:35:30.767494+01:00 ;  Date=2020-09-21T07:26:10+00:00 ;  Date=2020-09-21T07:26:10 ; Date=2020-09-21 ; |
| **N**_**spartitions =** 3:CO1_ABGD, 0.98 **/** test_BPP, 0.95 **/** PCA_phenotype,?  ;  *The spartition scores can also be omitted ( e.g. if no score at all), in which case the command would read:*  **N**_**spartitions =** 3:CO1_ABGD **/** test_BPP **/** PCA_phenotype; | *Number of species partitions; list of spartitions names ,* ***spartition score (? If no score)***  *Spartition names are separated by slashes.*  *This command define the order the spartitions (1^st^=CO1, 2^nd^=BPP, 3^rd^=PCA) that will be reused in the subsequent commands.*  *Two different spartitions are not allowed to share the same name*  *[note: spartition scores are included with number and names of spartitions and not in an optional additional command to facilitate extraction of information by human readers]* |
| **N_individuals =** 5 **/** 5 **/** 4; | *Total number of individuals (=samples, specimens) (1^st^, 2^nd^ and 3^rd^ spartition)*  *(the spartition order is the same as in N_spartitions)* |
| **N**_**subsets =** 3**:**0.95**,**0.98**,**0.99 **/** 2:0.95,0.98 **/** 4:?,?,?,?;  *The subset scores can also be omitted, in which case the command would read:*  **N**_**subsets =** 3 **/** 2 **/** 4; | *Total number of delimited subsets (1^st^, 2^nd^ and 3^rd^ spartition), subset score (? If no scores)*  *[note 1: The first score corresponds to the first subset appearing in the Individual assignment list (from top to bottom), the second subset score correspond to the second subset appearing in the list, etc.*  *Therefore, it is a “top to bottom” order based on individual assignment list, independently from the “value” of the number used to assign an individual to a given subset.*  *[note 2: subset scores are included after the number of subsets ]* |
| **[**CO1_ABGD : this is my first comment**]**  **[**CO1_ABGD : this is my second comment**]** **[**PCA_phenotype : this is  my first comment  extracted from the  third method  **]**  **[**my_three_delimitations : possible comment related to the concatenated multiple partition file **]** | *[Comment]*  *A comment begins with an opening bracket and finish with a closing bracket.*  *A multiple (concatenated) species partition file is able to report all the comments made independently in the different single species partition file (SPF), so the name of each SPF should be reported at the beginning of each comment.*  *[note: A comment, if needed, can be place anywhere in the file (see below). It can be in a single line (ex. both comment of CO1_ABGD) or on different lines (ex. PCA_Phenotype)]* |
| **Individual_assignment =**  Drosophila_32 **:** 1 **/** 1 **/** 4 **[**a comment can be placed anywhere**]**  Sample_2 **:** 1 **/** 1 **/** 3  Drosophila_China **:** 2 **/**2 **/** 2  Sample_E554 **:** 2 **/** 1 **/** ?  Droso_Vietnam **:** 3 **/** 2 **/** 1;  **[**CO1-ABGD : comment about the CO1 assignment**]**  **[**PCA-ABGD : comment about the PCA assignment**]** | *List of individuals (samples) with their respective assignment in each of the three spartitions (1^st^, 2^nd^ , then 3th spartitions), i.e., usually by each of three methods (? If individual not assigned by one of these methods).*  *Two different samples are not allowed to share the same name.*  *End of lines are separating each individual line :*  Drosophila_32**:** 1 **/** 1 **/** 4 ⏎  Sample_2 **:** 1 **/** 1 **/** 3 ⏎  Drosophila_China**:** 2 **/**2 **/** 2 ⏎  Sample_E554**:** 2 **/** 1 **/** ? ⏎  Droso_Vietnam**:** 3 **/** 2 **/** 1; |
| **end;** | *Ending line of the spart file. Indicates the end of a block. (the entire spart file is conceived as a single block). This line must be at the every end of the spart file (i.e., after the very last included command (wether they are compulsory or optional).* |
| **Optional commands** | *These optional fields are not part of the basic, compulsory spart syntax. They may be present in the spart files (and will be carried over or specifically generated if various spart files are merged into one), but* ***if they are missing it should not generate an error message****, except in such programs that specifically expect/require the information of some of these optional fields.*  *Remember: In general terms, the spart readers/parsers/analyzers should work in a way that they simply ignore lines with information they do not "understand" so that it becomes easy to add additional optional fields if it is later deemed to be useful for some specific applications.* |
| **Individual_score =**  Drosophila_32**:** ? **/** 0.99 **/**1.00  Sample_2 **:** ? **/**? **/** 1.00  Drosophila_China**:** ? **/**0.97 **/** 0.99  Sample_E554**:** ? **/**0.85 **/** ?  Droso_Vietnam**:** ? **/**0.99 **/** 0.96; | *List of individuals (samples) with their respective individual* ***score*** *according to each method, ie. 1st, 2^nd^ then 3^rd^ spartitions (? if no score).*  *Optional command: no need to present this command, e.g. if it is totally without values.*  *End of lines are separating each individual line :*  **Individual_score =**  Drosophila_32**:** ? **/** 0.99 **/**1.00 ⏎  Sample_2 **:** ? **/**? **/** 1.00⏎  Drosophila_China**:** ? **/**0.97 **/** 0.99⏎  Sample_E554**:** ? **/**0.85 **/** ? ⏎  Droso_Vietnam**:** ? **/**0.99 **/** 0.96;  It is important to accept either Windows (CR-LF), Unix (LF) and old Mac (CR) end of line  *[note: individual scores are given as separate (optional) command and are not included in the Assignment command because often they will be missing altogether, and because presenting them separately facilitates extraction of information in the Assignment command by human readers]* |
| **Spartition_score_type =** likelihood / ? /  ?;  **Subset_score_type =** bootstrap / ? / posterior_probability;  **Individual_score_type =** probability / bootstrap / ?; | *If these commands are absent, then the respective score types are missing = "?"*  *[note: Score types are flexibles (no defined list of type)]* |
| **Tree =**  test_BPP : ((Drosophila_32, Sample_2), Drosophila_China,(Sample_E554, Droso_Vietnam))  CO1_ABGD : ((Drosophila_China, Sample_2), Drosophila_32,(Sample_E554, Droso_Vietnam))  ; | *Reports (chain of characters) the input tree used for the calculation of a certain partition, in Newick format.*  *Multiple trees can be included, especially in a multi-spart file (one or maybe even several for each partition)* |
| **Command_line =**  test_BPP : 2019-01-30T09:26:10+00:00 / BPP version 2.0 / "bla bla bla bla bla bla bla bla bla bla bla bla bla bla bla bla bla bla bla bla bla bla bla bla bla bla bla bla bla bla bla bla bla bla bla bla bla bla bla bla bla bla bla bla bla bla bla bla bla bla bla bla bla bla bla bla”  CO1_ABGD : 2019-01-30T09:26:10+00:00 / 3.0 available at abgd.com / "abgd myinputfile.fas -a -v -P 0.3"  ; | *Gives the full command line of the program that was executed for generating the delimitation for the respective species partition (with the date, if existing) ).*  ***Commandline*** ***space*** *=*  ***name of the respective species partition*** *:* ***date of the original analysis*** */* ***version of the tool used*** */* ***specific commandline that was executed*** *(in quotation marks)*  *Question marks in case of (partly) missing data.*  ***semicolon*** *(to end the line).* |

**5. Format description: SPART.XML**

This format follows the Extensible Markup Language (XML), a markup language that defines a set of rules for encoding documents in a format that is both human-readable and machine-readable.

The spart.xml format encodes the same information as the matricial spart, but with a vocabulary adapted to fit conventions and requirements of XML. In particular this affects the following commands:
spartition_score = spartitionScore
individual_score = individualScore
individual_score_type = individualScoreType
subset_score = subsetScore

spartition_score_type and subset_score_type are not used as separate terms, but the respective information encoded in the respective lines subsetScore and and spartitionScore under "type"; the respective values are given in the same lines under "value". Subset scores are furthermore placed in a section "external support" which can also provide information ("source") on the type of analysis, algorithm or program this support was derived from.

```
<?xml version="1.0" ?>
<root>
	<project_name>Mantella.fas</project_name>
	<date>2021-01-29T18:13:35</date>
    <!-- Generated by bPTP -->
    <!-- WARNING: The sample names below may have been changed to fit SPART specification (only alphanumeric characters and _ ) -->
    <!-- user comment: this analysis was generated based on a single ML tree obtained in MEGA 7 -->
```

<individuals>	*List of individuals sampled*

```
        <individual id="aura_ZCMV1234"  />
        <individual id="aura_ZCMV1235"  />
        <individual id="aura_ZCMV1236"  />
        <individual id="aura_ZCMV1237"  />
        <individual id="aura_ZCMV1238"  />
        <individual id="aura_ZCMV1239"  />
        <individual id="aura_FGZC987"   />
        <individual id="aura_FGZC986"   />
        <individual id="crocea_ZCMV234" />
        <individual id="crocea_ZCMV235" />
        <individual id="miloty_ACZC324" />
        <individual id="miloty_ACZC329" />
        <individual id="crocea_ZCMV236" />
        <individual id="crocea_ZCMV237" />
        <individual id="miloty_ACZV679" />
        <individual id="miloty_ZCMV479" />
	</individuals>
```

<spartitions>	*Spartitions description*

<spartition label="Mantella_bPTP" spartitionScore="1.234E-6" spartitionScoreType="logLikelihood" > *First spartition description*

<remarks>First spartition</remarks>

<subsets>	*Subsets description of the first spartition (N=3 in this exemple)*

```
				<subset label="1">
				    <externalSupport>
				        <subsetScore type="posterior" value="1.23E-6" source="BEAST analysis 2021-03-02" />
                                          </externalSupport>
					<individual ref="aura_ZCMV1234" individualScore="1.23E-3" individualScoreType="probability" />
					<individual ref="aura_ZCMV1235" individualScore="1.23E-3" individualScoreType="probability" />
					<individual ref="aura_ZCMV1236" individualScore="1.23E-3" individualScoreType="probability" />
					<individual ref="aura_ZCMV1237" individualScore="1.23E-3" individualScoreType="probability" />
					<individual ref="aura_ZCMV1238" individualScore="1.23E-3" individualScoreType="probability" />
					<individual ref="aura_ZCMV1239" individualScore="1.23E-3" individualScoreType="probability" />
					<individual ref="aura_FGZC987"  individualScore="1.23E-3" individualScoreType="probability" />
					<individual ref="aura_FGZC986"  individualScore="1.23E-3" individualScoreType="probability" />
				</subset>
				<subset label="2">
				    <externalSupport>
				        <subsetScore type="posterior" value="7.34E-6" source="BEAST analysis 2021-03-02" />
                                           </externalSupport>
					<individual ref="crocea_ZCMV234" individualScore="1.23E-3" individualScoreType="probability" />
					<individual ref="crocea_ZCMV235" individualScore="1.23E-3" individualScoreType="probability" />
					<individual ref="miloty_ACZC324" individualScore="1.23E-3" individualScoreType="probability" />
					<individual ref="miloty_ACZC329" individualScore="1.23E-3" individualScoreType="probability" />
					<individual ref="crocea_ZCMV236" individualScore="1.23E-3" individualScoreType="probability" />
					<individual ref="crocea_ZCMV237" individualScore="1.23E-3" individualScoreType="probability" />
					<individual ref="miloty_ACZV679" individualScore="1.23E-3" individualScoreType="probability" />
					<individual ref="miloty_ZCMV479" individualScore="1.23E-3" individualScoreType="probability" />
				</subset>
				<subset label="3">
				    <externalSupport>
				        <subsetScore type="posterior" value="1.01E-5" source="BEAST analysis 2021-03-02" />
                                          </externalSupport>
					<individual ref="miloty_ACZV679" individualScore="1.23E-3" individualScoreType="probability" />
					<individual ref="miloty_ZCMV479" individualScore="1.23E-3" individualScoreType="probability" />
				</subset>
			</subsets>
		</spartition>
```

<spartition label="analysis P2"> <score type="likelihood" value="1.0345E-06" /> *Second spartition description*

<remarks>Second spartition</remarks> *Subsets description of the 2^nd^ spartition (N=5 in this exemple)*

```
			<subsets>
				<subset label="1">
					<individual ref="aura_ZCMV1234" />
					<individual ref="aura_ZCMV1235" />
					<individual ref="aura_ZCMV1236" />
					<individual ref="aura_ZCMV1237" />
				</subset>
				<subset label="2">
					<individual ref="aura_ZCMV1238" />
					<individual ref="aura_ZCMV1239" />
				</subset>
				<subset label="3">
					<individual ref="aura_FGZC987" />
					<individual ref="aura_FGZC986" />
				</subset>
				<subset label="4">
					<individual ref="crocea_ZCMV234" />
					<individual ref="crocea_ZCMV235" />
					<individual ref="miloty_ACZC324" />
					<individual ref="miloty_ACZC329" />
					<individual ref="crocea_ZCMV236" />
					<individual ref="crocea_ZCMV237" />
					<individual ref="miloty_ACZV679" />
					<individual ref="miloty_ZCMV479" />
				</subset>
				<subset label="5">
					<individual ref="miloty_ACZV679" />
					<individual ref="miloty_ZCMV479" />
				</subset>
			</subsets>
		</spartition>
	</spartitions>
</root>
```
